# TS-AI: A deep learning pipeline for multimodal subject-specific parcellation with task contrasts synthesis

**DOI:** 10.1101/2024.06.14.598994

**Authors:** Chengyi Li, Yuheng Lu, Shan Yu, Yue Cui

**Author notes:** Corresponding authors: Shan Yu, Institute of Automation, Chinese Academy of Sciences, Beijing 100190, China, and Yue Cui, Institute of Automation, Chinese Academy of Sciences, Beijing 100190, China.

## Abstract

Accurate mapping of brain functional subregions at an individual level is crucial. Task-based functional MRI (tfMRI) captures subject-specific activation patterns during various functions and behaviors, facilitating the individual localization of functionally distinct subregions. However, acquiring high-quality tfMRI is time-consuming and resource-intensive in both scientific and clinical settings. The present study proposes a two-stage network model, TS-AI, to individualize an atlas on cortical surfaces through the prediction of tfMRI data. TS-AI first synthesizes a battery of task contrast maps for each individual by leveraging tract-wise anatomical connectivity and resting-state networks. These synthesized maps, along with feature maps of tract-wise anatomical connectivity and resting-state networks, are then fed into an end-to-end deep neural network to individualize an atlas. TS-AI enables the synthesized task contrast maps to be used in individual parcellation without the acquisition of actual task fMRI scans. In addition, a novel feature consistency loss is designed to assign vertices with similar features to the same parcel, which increases individual specificity and mitigates overfitting risks caused by the absence of individual parcellation ground truth. The individualized parcellations were validated by assessing test-retest reliability, homogeneity, and cognitive behavior prediction using diverse reference atlases and datasets, demonstrating the superior performance and generalizability of TS-AI. Sensitivity analysis yielded insights into region-specific features influencing individual variation in functional regionalization. In addition, TS-AI identified accelerated shrinkage in the medial temporal and cingulate parcels during the progression of Alzheimer’s disease, suggesting its potential in clinical research and applications.

## 1. Introduction

The human brain exhibits significant individual variability in structural and functional organizations (Frost and Goebel, 2012; Mueller et al., 2013). Brain atlases, delineating anatomically distinct and functionally specialized cortical subregions, serve as valuable tools in neuroscience research and clinical practice (Fan et al., 2016; Glasser et al., 2016).

However, directly registering these atlases to individual spaces based on morphological information often overlooks inter-subject differences in regional positions and topography, leading to inaccuracies in individual atlas mapping (Glasser et al., 2016). Consequently, accurate mapping of functional regions at the individual level is pivotal for a comprehensive understanding of the variations in brain function and behavior (Keller et al., 2024; Shanmugan et al., 2021; Zhou et al., 2023), enabling precise diagnosis of brain disorders (Meng et al., 2021; Zhao et al., 2023), as well as personalizing treatments such as targeting in transcranial magnetic stimulation (Ren et al., 2023) and deep brain stimulation (Patriat et al., 2018).

Magnetic resonance imaging (MRI) is a non-invasive technique for measuring brain structure and function, and is essential for obtaining individual-specific brain mapping. Researchers have developed individual-specific atlases using resting-state functional MRI (rsfMRI) (Gordon et al., 2017c; Honnorat et al., 2017; Kong et al., 2019; Li et al., 2023; Wang et al., 2015), diffusion MRI (dMRI) (Han et al., 2020; Ma et al., 2022), structural MRI (sMRI) (Bayrak et al., 2022), and task-based functional MRI (tfMRI) (Glasser et al., 2016). Evidence has shown that integrating multiple modalities provides comprehensive profiles of brain function and structure compared to using single-modality MRI data (Glasser et al., 2016; Parisot et al., 2017; Wang et al., 2018). For instance, the combination of dMRI, rsfMRI, and myelin maps for individual parcellation using Markov random fields has shown improved alignment with myelin and cytoarchitectural maps compared to single-modality data (Parisot et al., 2017). Similarly, Wang et al. (2018) demonstrated that combining local anatomical and functional connectivity data derived from dMRI and rsfMRI through clustering enhances the test-retest reliability of individual parcellations compared to single-modality alternatives. Glasser et al. (2016) identified individual regions using binary classifiers based on rsfMRI, tfMRI, visuotopic and morphological data. It is found that tfMRI is particularly crucial for capturing individual activation patterns across a broad spectrum of functions and behaviors, from primary sensorimotor to higher-order cognitive and emotional processes, facilitating the localization of functionally distinct subregions at an individual level. However, acquiring a battery of tfMRI data is time-consuming and resource-intensive, and high-quality tfMRI tasks are difficult to complete for many participants, especially patients with functional impairments, the elderly, and children, limiting its applicability in scientific and clinical settings. Studies have shown that tfMRI-based functional activations can be predicted by anatomical connectivity (Osher et al., 2016; Saygin et al., 2012; Saygin et al., 2016; Wu et al., 2020), as well as functional networks and connectivity (Bernstein-Eliav and Tavor, 2022; Ngo et al., 2022; Tavor et al., 2016; Tik et al., 2023; Tobyne et al., 2018; Zheng et al., 2022), ranging from primary somatosensory to high-order association cortices. Moreover, the synthesized functional activations can be utilized in downstream applications, such as the prediction of general cognitive ability (Gal et al., 2022). Therefore, synthesizing functional activations and integrating them with other MRI modalities is a promising approach for accurate localization of subject-specialized brain regions.

There has been a recent interest in employing deep learning-based methods for individualizing parcellation (Glasser et al., 2016; Li et al., 2023; Ma et al., 2022; Williams et al., 2021; Zhao et al., 2018), which involves estimating a model for mapping the individual imaging signals to individualized parcellations. These deep learning-based methods differ from non-deep learning approaches, such as clustering (Wang et al., 2015), template matching (Gordon et al., 2017a; Gordon et al., 2017b), and graph partition methods (Chong et al., 2017; Cui et al., 2024; Honnorat et al., 2017), which directly derive individual parcellations from individual brain imaging signals with the constraints of prior assumptions. In contrast, deep learning-based approaches can automatically learn high-order and nonlinear relationships from brain features to subregional specialization at an individual level, potentially better reflecting the anatomical and functional complexity of the brain. Nonetheless, a major challenge in deep learning-based methods is the lack of definitive individual parcellation as the ground truth to supervise the model for automated learning of individual parcels. To mitigate this issue, some research has used alternative individualization methods such as Glasser’s private individualized atlases (Williams et al., 2021), Kong’s Multi-session hierarchical Bayesian model (Bayrak et al., 2022), or matrix decomposition (Zhao et al., 2018) to generate proxy ground truth for training their models. Other studies have employed parametric low-dimensional models with referring group-level atlases as ground truth (Glasser et al., 2016; Ma et al., 2022; Qiu et al., 2022). For instance, Glasser et al. (2016) used a three-layer multilayer perceptron (MLP) to determine if a vertex belongs to a specific parcel, with a sufficient number of samples to mitigate overfitting risks. However, individual parcellations may be affected by some imprecise labels from either reference or individualized atlases that serve as the only constraint in the learning procedure. To counteract the absence of individual atlas ground truth, incorporating constraints like within-parcel homogeneity into the model could be a preferable approach.

In the present study, we propose a two-stage network framework, termed Atlas Individualizing with Task contrasts Synthesis (TS-AI), for obtaining subject-specific atlases across the cortex. In the first stage, the model synthesizes task contrast maps for each individual, incorporating tract-wise anatomical connectivity and resting-state networks. In the second stage, these synthesized maps, along with feature maps of tract-wise anatomical connectivity and resting-state networks, are fed into an end-to-end deep neural network for atlas individualization. The advantage of the present study includes the synthesized task contrast maps predicted from rsfMRI and dMRI for individual parcellation without the acquisition of actual task fMRI scans. Another contribution is a novel feature consistency loss which increases individual specificity by assigning vertices with similar features to the same parcel. This is effective in dealing with the lack of individual atlas ground truth, enabling the sufficient utilization of deep networks’ capabilities in adaptive feature extraction and end-to-end learning for atlas individualization. The supervised atlas consistency loss, which ensures the similarity between individualized and reference atlas, is incorporated with the unsupervised feature consistency loss as the loss function of the second stage. Note that no individual parcellations are used as supervisory information in the learning process of TS-AI. Comprehensive validations including test-retest reliability, homogeneity analysis, and behavioral prediction were conducted using Human Connectome Project (HCP) and independent Chinese HCP (CHCP) datasets. Additionally, we performed sensitivity analyses to evaluate the contributions of input features on the individualization of each functional region. Furthermore, we applied TS-AI to an Alzheimer’s disease cohort, and observed that the areal variations of TS-AI-derived regions reflect the progression of AD, suggesting its potential for monitoring disease progression in neuropsychiatric disorders.

## 2. Materials and Methods

### 2.1. Datasets acquisition, MRI preprocessing, and features extraction

#### 2.1.1. HCP dataset

The present study utilized the Human Connectome Project (HCP) for training, inference, and validation of the TS-AI model. The T1- and T2-weighted structural MRI, rsfMRI, tfMRI, and dMRI data were obtained from the HCP S1200 release. These images were preprocessed using the HCP minimal preprocessing pipelines (https://github.com/Washington-University/HCPpipelines). From the acquired data, we selected a subset of 968 participants (454 males, aged 22 ∼ 37 years) who had successfully preprocessed multimodal images for further analysis. The study also incorporated behavioral data (n = 953) and retest imaging data (n = 39).

The T1- and T2-weighted structural MRI data underwent cortical surface reconstruction using the HCP preprocessing pipeline. Nonlinear surface registrations from individual surfaces to the template were performed using MSM-Sulc and MSM-All (Robinson et al., 2014) to ensure accurate alignment of surfaces. The rsfMRI data had undergone the minimal fMRI preprocessing employed by HCP, including motion correction, slice timing correction, susceptibility-induced distortion correction, functional-structural registration, and independent component analysis (ICA)-based artifact removal. Additionally, temporal band-pass filtering was applied to retain frequencies within the 0.01 to 0.1 Hz range, and spatial smoothing was conducted using a 4 mm Gaussian filter with a full width at half maximum (FWHM) on the mid-thickness surface. Group ICA with 100 dimensions was performed using MELODIC from FSL (FMRIB Software Library) by HCP, and we selected 47 non-noise independent components on the cortical surface. We selected a relatively high-dimensional ICA because such dimensionality provides refined networks that correspond well to known anatomical and functional specializations (Allen et al., 2011; Salman et al., 2019). Individual resting-state functional networks were then extracted via group information guided ICA (GIG-ICA) (Du and Fan, 2013). This process projected group independent components onto individual participants, driven by their specific resting-state time series. As a result, the rsfMRI-derived features comprised 47 individualized resting-state networks.

The tfMRI data had undergone the minimal fMRI preprocessing and tfMRI analysis conducted by HCP, which included seven task domains: emotion, gambling, language, motor, relational, social, and working memory, each comprising multiple contrasts. To reduce redundancy among these contrasts, we excluded contrasts between conditions, negative contrasts, and average contrasts, resulting in 24 task activation contrast maps that focused on events alone (Ito et al., 2020). A list of these task contrasts can be found in Supplementary Table 1.

The dMRI data had undergone denoising, eddy correction, susceptibility-induced distortion correction, diffusional-structural registration, and fiber orientation estimation (FSL’s bedpostX) in HCP release. We then conducted whole-brain tractography for each hemisphere separately, seeding 5000 streamlines from each vertex on the white matter surface. These streamlines were projected onto whole-brain voxels with 5 mm isotropic resolution and subsequently log-transformed. This process yielded a V-by-W voxel-wise anatomical connectivity profile, where V is the number of cortical vertices and W is the number of whole-brain voxels, for each participant. Additionally, 72 individual fiber tract maps were generated using TractSeg (Wasserthal et al., 2018). Each map was normalized to ensure an equal sum (set to be 1). Tract-wise anatomical connectivity was then calculated by summing the connectivity profile values within the corresponding tract, formulated as 𝐴 = 𝑇𝑃, where 𝐴 denoted a B-by-V matrix as tract-wise anatomical connectivity, 𝑇 denoted a B-by-W matrix as normalized tract maps, 𝑃 denoted a W-by-V matrix as voxel-wise anatomical connectivity, and B = 72 denoted the number of tracts. The anatomical connectivity features used in TS-AI were then extracted from 𝐴, resulting in 41 maps for each hemisphere, where the tracts from its opposite hemisphere were aborted.

#### 2.1.2. CHCP dataset

The Chinese Human Connectome Project (CHCP) served as an independent validation dataset for analyzing the homogeneity of anatomical and resting state connectivity, and task activations. The original T1- and T2-weighted structural MRI, rsfMRI, tfMRI, and dMRI data for 366 participants were downloaded from CHCP (http://chinese-hcp.cn/) and obtained under informed consent. The preprocessing of these data was identical to that of the HCP dataset, including the extraction of resting-state functional, task functional, and anatomical connectivity features. Only participants with complete and successfully processed data across all modalities were included in further analyses, resulting in a final cohort of 315 participants (145 males, aged 18 ∼ 79 years). CHCP participants can be grouped into young (n = 221, aged 18 ∼ 33 years) and old (n = 94, aged 48 ∼ 79 years) groups.

#### 2.1.3. ADNI dataset

Alzheimer’s Disease Neuroimaging Initiative (ADNI) provided an independent dataset used for validating the efficacy of individualized atlases as potential markers in Alzheimer’s disease (AD) progression. We included 252 participants from the ADNI-3 cohort, who have undergone T1- and T2-weighted structural MRI, rsfMRI, and dMRI scans, encompassing 56 AD patients, 98 with mild cognitive impairment (MCI), and 98 age- and gender-matched healthy controls (HC). Informed consent was obtained from all participants.

The preprocessing of the structural MRI data is identical to that of HCP dataset. For rsfMRI, our preprocessing steps included slicing timing correction, motion correction, functional-structural registration, temporal band-pass filtering (0.01 ∼ 0.1 Hz), and spatial smoothing (4 mm FWHM Gaussian filter) on the mid-thickness surface (https://github.com/ThomasYeoLab/Standalone_CBIG_fMRI_Preproc2016). The dMRI data preprocessing included denoising, eddy correction, diffusional-anatomical registration, fiber orientation estimation, whole-brain tractography, and fiber tract segmentation, consistent with the protocols employed on the HCP and CHCP datasets. Subsequently, features including resting-state networks and anatomical connectivity were extracted. After preprocessing, 10 (5 AD, 1 MCI, and 4 HC), 5 (2 AD, 2 MCI, and 1 HC) and 9 (5 AD, 1 MCI, and 3 HC) participants were excluded due to failed surface construction, low-quality rsfMRI and low-quality dMRI, respectively. Consequently, the final subset comprised 44 AD patients (age range 55 ∼ 87, 75.2 ± 7.71 years), 94 MCI participants (age range 49 ∼ 92, 74.0 ± 7.76 years), and 90 HC participants (age range 57 ∼ 95, 72.7 ± 8.85 years) with successfully processed data across all modalities.

The demographic characteristics of the participants and MRI resolutions of HCP, CHCP, and ADNI datasets are shown in Table 1 and Supplementary Table 2.

**Table 1.** Demographic characteristics of HCP, CHCP, and ADNI used in this study.

| Dataset | | Number of participants | Age range in years (mean ± SD) | Sex (Female/Male) |
| --- | --- | --- | --- | --- |
| HCP | | 968 | 22 ~ 37 ($28.7 \pm 3.7$) | 514/454 |
| CHCP | Young | 221 | 18 ~ 33 ($22.4 \pm 2.9$) | 111/110 |
| | Old | 94 | 48 ~ 79 ($62.6 \pm 5.5$) | 59/35 |
| | Total | 315 | 18 ~ 79 ($34.4 \pm 18.8$) | 170/145 |
| ADNI | HC | 90 | 57 ~ 95 ($72.7 \pm 8.9$) | 42/48 |
| | MCI | 94 | 49 ~ 92 ($74.0 \pm 7.8$) | 48/46 |
| | AD | 44 | 55 ~ 87 ($75.2 \pm 7.7$) | 28/16 |
| | Total | 228 | 49 ~ 95 ($73.6 \pm 8.3$) | 118/110 |

AD, Alzheimer's disease; ADNI, Alzheimer's Disease Neuroimaging Initiative dataset; CHCP, Chinese Human Connectome Project dataset; HC, healthy control; HCP, Human Connectome Project dataset; MCI, Mild Cognitive Impairment.

### 2.2. Atlas individualizing with task contrasts synthesis (TS-AI)

The proposed TS-AI comprises task contrasts synthesis and atlas individualizing sub-models. TS-AI is trained in two sequential stages. During the task contrasts synthesis stage, tract-wise anatomical connectivity and resting-state networks serve as inputs to generate task contrasts (Figure 1a). Subsequently, in the atlas individualizing stage, the synthesized task contrasts, along with the tract-wise anatomical connectivity, and resting-state networks, are used to train the atlas individualizing sub-model, ultimately producing individualized atlases (Figure 1b).

**Figure 1.**
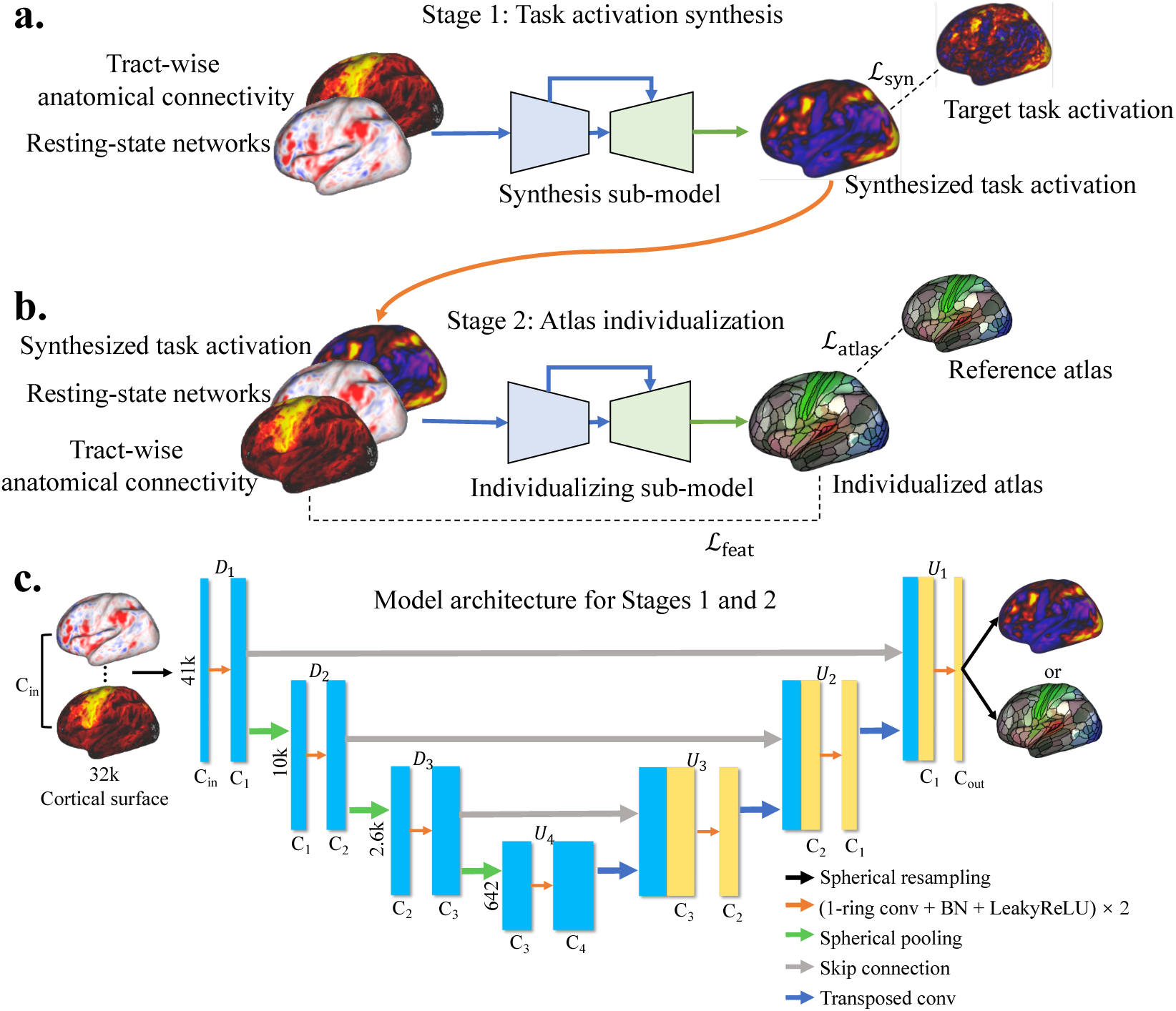
Schematic of the TS-AI (atlas individualizing with task contrasts synthesis) workflow. TS-AI consists of two stages, each corresponding to a sub-model: task contrasts synthesis (a) and atlas individualizing (b). **a.** In the task contrasts synthesis stage, the tract-wise anatomical connectivity, and resting-state networks are served as inputs to synthesize a battery of task contrasts. **b.** In the atlas individualizing stage, the synthesized task contrasts, along with tract-wise anatomical connectivity and resting-state networks, contribute to the generation of individualized atlases. **c.** The architecture for both sub-models is a Spherical U-Net, featuring three downsampling and four upsampling blocks. ℒ_syn_, task contrast map synthesis loss; ℒ_atlas_, atlas consistency loss; ℒ_feat_, feature consistency loss; conv, spherical convolution; BN, BatchNorm layer.

TS-AI utilizes the Spherical U-Net architecture (Zhao et al., 2019) for both the synthesis and individualizing sub-models. Spherical U-Net is characterized by its U-Net architecture and spherical convolution, operating on the cortical surface mesh (Figure 1c). The standard sphere for cortical surface representation in Spherical U-Net is derived from a regular icosahedron with 12 vertices. This is achieved by iteratively adding a new vertex to the center of each edge in each triangle, resulting in a series of icospheric meshes from the 0th to the 6th order, with the number of vertices increasing from 12 to 40,962. The encoder consists of 𝑀 − 1 downsampling blocks that progressively reduce the mesh resolution, while the decoder comprises 𝑀 upsampling blocks that gradually increase the mesh resolution. Furthermore, 𝑀 − 1 skip connections are established between the corresponding downsampling and upsampling blocks. In the current implementation, 𝑀 is set to 4, handling the 6th to 3rd order meshes in the 1st to 4th downsampling/upsampling blocks. Each block incorporates two spherical convolution layers (with channels of 64, 128, 256, and 256 for the 1st to 4th blocks, respectively), followed by BatchNorm and LeakyReLU activation, and an additional downpooling/uppooling layer when necessary. Each sub-model processes one hemisphere separately.

In both stages, the input features are z-score normalized across vertices within each feature map. Given that all features and atlases for each hemisphere are initially projected onto an fsLR 32k surface mesh (32,492 vertices), input features are resampled to the 6th order icospheric mesh (40,962 vertices) using barycentric interpolation implemented by “wb_command -metric-resample” from Connectome Workbench (https://humanconnectome.org/software/connectome-workbench). After processing, the output features and atlases are resampled back to the fsLR 32k surface mesh.

#### 2.2.1. Task contrasts synthesis

The synthesis sub-model employs tract-wise anatomical connectivity and resting-state networks to predict the task contrast maps. The loss function for this sub-model is defined as the mean squared error (MSE) between the synthesized and the target (measured) task contrast maps:

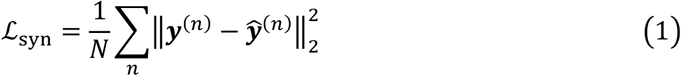

Here, 𝑁 represents the number of training samples, **y**^(n)^ represents the target task contrast for participant 𝑛, and ***ŷ***^(n)^ represents the synthesized task contrast.

#### 2.2.2. Atlas individualizing

In the atlas individualizing stage, the individualizing sub-model receives input from a fusion of synthesized task contrast, tract-wise anatomical connectivity, and resting-state network maps. This sub-model is designed to generate individualized atlases, guided by both feature consistency loss and atlas consistency loss, using one model for the individualization of all parcels.

Data augmentation is applied when training the individualizing sub-model. Specifically, for each participant, spherical nonlinear warping is applied to the input features and the reference atlas inspired by Williams et al. (2021) and Fawaz et al. (2021). The augmentation process involves random displacement of 100 vertices on the 2nd order icospheric mesh, with each displacement vector uniformly sampled from a ball space of 32 mm radius. The displaced vertices are then reprojected back onto the sphere, and these deformations are sequentially interpolated to 3rd, 4th, 5th, and 6th order icospheric meshes.

The loss function for the individualizing sub-model is formulated as ℒ_indiv_ = ℒ_atlas_ + λℒ_feat_, where ℒ_feat_ denotes feature consistency loss, ℒ_atlas_ denotes atlas consistency loss, and λ is a hyperparameter. The feature consistency loss aims to maximize the feature similarity within each parcel of the individualized atlas, calculated as the MSE between the input multimodal features (anatomical, resting-state, and task features) of each vertex and their average features within the corresponding parcel:

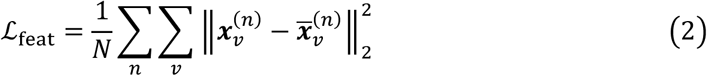

Here, 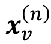 represents the input features at vertex 𝑣, and 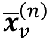 indicates the average input features within the parcel to which vertex 𝑣 belongs, which is computed as follows:

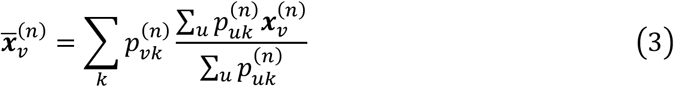

Here, 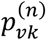 denotes the probability predicted by the sub-model that vertex 𝑣 belongs to parcel 𝑘, and 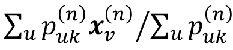 denotes the average input features of parcel 𝑘.

The atlas consistency loss is designed to constrain the differences between the individualized and reference atlases:

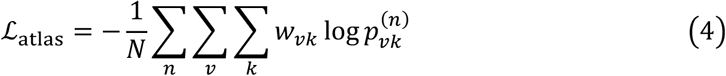

Here, 𝑤_01_ ∈ {0,1} indicates whether vertex 𝑣 belongs to parcel 𝑘 in the reference atlas. Notably, all training losses including task contrasts synthesis loss, feature consistency loss, and atlas consistency loss are computed based on the 40k sphere.

During model inference, the output probability 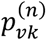 is converted to labels (i.e., individualized atlas) according to the winner-take-all principle. The resulting atlases then undergo postprocessing, as described by Glasser et al. (2016), which involves several steps: (i) removing very small patches smaller than 25 mm², (ii) merging fragmented patches separated by 3 or fewer vertices, (iii) eliminating patches smaller than 0.33 times the size of the largest patch, (iv) removing patches over 30 mm from their nearest patch within the same parcel, and (v) assigning unlabeled vertices, removed in (i), (iii), and (iv), to the nearest parcel with labeled vertices.

### 2.3. Experimental setup

The synthesis sub-model was trained for 30 epochs with a batch size of 8, using the Adam optimizer set at a learning rate of 10^-3^. The training was completed in approximately one hour on a single NVIDIA RTX3090 GPU. An early-stopping strategy was employed, selecting the final model based on the highest Pearson’s correlation coefficient between the synthesized and target task contrasts.

For the individualizing sub-model, training was conducted for 30 epochs with a batch size of 4, also employing the Adam optimizer with a learning rate of 10^-3^. The hyperparameter λ was set to be 10 (see Supplementary Table 3 for evaluations of different 𝜆). This training process took around 1.5 hours, also on a single NVIDIA RTX3090 GPU. Reference parcellations included the Glasser (Glasser et al., 2016) and Brainnetome (Fan et al., 2016) atlases, comprising 180 and 105 cortical parcels for each hemisphere, respectively. The optimal model was chosen based on the lowest feature consistency loss during the entire training phase.

Model training and evaluations were carried out on HCP dataset using a 2-fold cross-validation. A detailed description of the cross-validation can be found in Supplementary Methods and Supplementary Figure 1. Participants were randomly divided into two folds (n = 484). Within each fold, 80% of participants (n = 387) were used for training, and the remaining 20% (n = 97) were used for model selection. The entire cohort of the other fold was used for evaluation. The better model trained from the 2-fold cross-validation was subsequently applied to the CHCP and ADNI datasets for atlas individualization and further analysis.

### 2.4. Evaluations for task contrasts synthesis

The correlation between predicted and target contrast maps was quantitatively assessed using Pearson’s correlation coefficient and the Dice coefficient. The Dice coefficient measures the overlap between the predicted and target contrast maps within the activated vertices. Given a threshold 𝑡, the Dice coefficient is computed as:

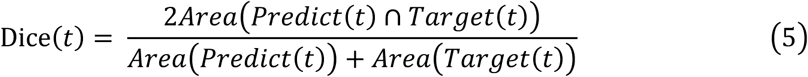

Here, 𝐴𝑟𝑒𝑎(𝑃𝑟𝑒𝑑𝑖𝑐𝑡(𝑡)) denotes the area of the top 𝑡 activated vertices in the predicted contrast map, 𝐴𝑟𝑒𝑎(𝑇𝑎𝑟𝑔𝑒𝑡(𝑡)) denotes the area of the top 𝑡 activated vertices in the target contrast map, and 𝐴𝑟𝑒𝑎(𝑃𝑟𝑒𝑑𝑖𝑐𝑡(𝑡) ∩ 𝑇𝑎𝑟𝑔𝑒𝑡(𝑡)) denotes the area of overlap between the predicted and target maps at the specified threshold. Further, the Dice AUC (area under the Dice curve) was employed as a measure of prediction quality on activated areas (Ngo et al., 2022). It is computed by integrating Dice(𝑡) from the threshold range of 5% to 50%, in increments of 0.01.

### 2.5. Evaluations for atlas individualization

To assess the individualization performance of TS-AI, we conducted comprehensive evaluations using both quantitative and qualitative metrics. TS-AI was benchmarked against individualized multimodal parcellation (IMMP), a learning-based multimodal individualizing method (Glasser et al., 2016). IMMP employed MLPs as binary classifiers for determining if a vertex belonged to a parcel in the reference atlas. Each MLP was structured with three layers: an input layer, a hidden layer with 9 nodes, and an output layer with 2 nodes. The hidden layer incorporated the ReLU activation function, whereas the output layer used the Sigmoid activation function. Additionally, cross-entropy loss was employed during the training phase. To ensure a fair comparison, the input features used for IMMP were identical to those of TS-AI.

#### 2.5.1. Test-retest reliability

The reliability of an individualized atlas was assessed by comparing intra-subject overlap rates (within the same individual) against inter-subject overlap rates (between different individuals). Higher intra-subject overlap rates compared to inter-subject overlap rates indicate that the individualized atlas successfully maintains consistency within each participant while capturing variability across different participants. Quantitatively, test-retest reliability can be measured by the effect size (Cohen’s d) of inter- and intra-subject overlap rates, which is defined as:

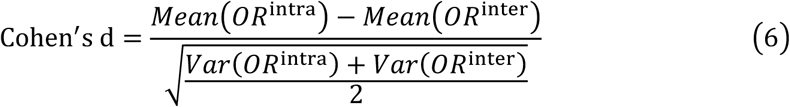

where 𝑀𝑒𝑎𝑛(𝑂𝑅^intra^) and 𝑉𝑎𝑟(𝑂𝑅^intra^) represent the mean and variance of the overlap rates within subjects, and 𝑀𝑒𝑎𝑛(𝑂𝑅^inter^) and 𝑉𝑎𝑟(𝑂𝑅^inter^) represent those metrics between subjects.

#### 2.5.2. Cognitive behavior prediction

Previous studies (Kong et al., 2019; Ma et al., 2022) have established that the topography of individual parcellations can predict cognitive behaviors using Kernel Ridge Regression (KRR). For each given participant 𝑗, KRR predicts the corresponding behavioral score 𝑠_5_based on the similarity between participant 𝑗’s individualized atlas and those of participants in the training set. The optimization objective can be formulated as:

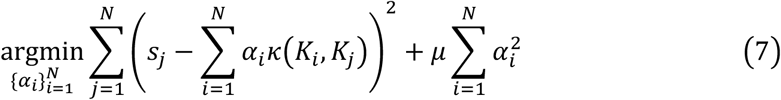

Here, 𝛼_i_ denotes the KRR parameter that signifies the contribution of the 𝑖-th participant in the training set, 𝜅(𝐾_i_, 𝐾_j_) denotes the overlap rate between individualized atlases 𝐾_i_ and 𝐾_j_, 𝑁 denotes the number of participants in the training set, and 𝜇 is a hyperparameter of the parameter regularization term, which is empirically set to 1.

Behavioral prediction was performed in the two HCP folds separately, each with 5-fold cross-validation for 100 repetitions to ensure robustness (Supplementary Methods and Supplementary Figure 1). The final prediction performance was determined as the average of Pearson’s correlation coefficients (PCC) between the predicted and measured behavioral scores across 1000 experiments (i.e., 5 × 100 × 2).

#### 2.5.3. Homogeneity

We employed three distinct homogeneity measures to assess the consistency within individualized parcels of the atlases: anatomical connectivity homogeneity (ACH), resting-state functional homogeneity (RFH), and task activation homogeneity (TAH), computed based on anatomical connectivity, functional connectivity, and task activation, respectively.

ACH is calculated using the average Pearson’s correlation coefficient of the dMRI-derived anatomical connectivity profiles between vertex pairs within each parcel (Han et al., 2020):

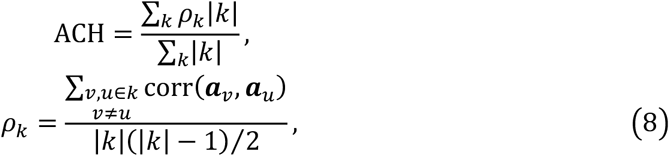

Here, corr(𝒂_0_, 𝒂_2_) denotes the Pearson’s correlation coefficient between the voxel-wise anatomical connectivity of vertices 𝑣 and 𝑢 within parcel 𝑘, and |𝑘| denotes the number of vertices in parcel 𝑘.

RFH measures the Pearson’s correlation coefficient of functional timeseries between vertex pairs within each parcel (Chong et al., 2017; Kong et al., 2019):

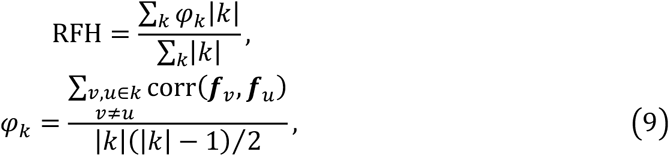

Here, corr(𝒇*_v_*, 𝒇*_u_*) denotes the Pearson’s correlation coefficient of the functional timeseries between vertices 𝑣 and 𝑢 within parcel 𝑘.

TAH is defined as the inverse standard deviation of the z-statistics of task activation within a parcel (Chong et al., 2017; Kong et al., 2019). For each task, the TAH of a task contrast for a subject is the average homogeneity weighted by the parcel’s area:

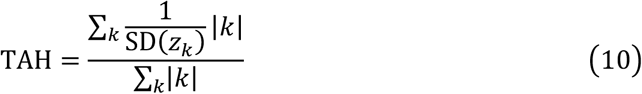

Here, SD(𝑧_1_) denotes the standard deviation of task activation z-statistics for parcel 𝑘.

To assess the generalizability of TS-AI, we applied the trained model to an independent dataset, CHCP, and compared ACH, RFH, and TAH in TS-AI-derived atlases with those from IMMP-derived and registration-based atlases. The calculations for ACH, RFH, and TAH in CHCP participants were identical to those used for HCP participants.

#### 2.5.4. Interpretability via sensitivity analysis

To explore the interpretability of features in the individualizing sub-model, we conducted both modality-wise and feature-wise sensitivity analyses. In the modality-wise sensitivity analysis, we set all input feature maps of each modality to 0 while maintaining the feature maps of other modalities. This perturbation resulted in modified individualized atlases. We then quantified the sensitivity of each parcel to a specific modality by defining an individual sensitivity measure as:

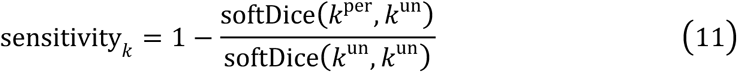

Here, 𝑘^per^denotes parcel 𝑘 in the perturbed atlas, and 𝑘^un^denotes parcel 𝑘 in the unperturbed atlas. The soft Dice coefficient is defined as:

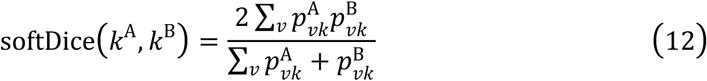

Here, 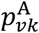 denotes the probability that vertex 𝑣 belongs to parcel 𝑘 in atlas A. We then averaged these individual sensitivity maps to create a group-level sensitivity map, where a higher value signified the higher sensitivity of a parcel to the specific modality. Similarly, for feature-wise sensitivity analysis, each input feature map was set to 0 while the other feature maps were retained. This process was repeated to finally yield a group-level sensitivity map for each feature.

### 2.6. TS-AI application

In order to assess its potential impact on neurological and psychiatric disease research, the TS-AI model, initially trained on the healthy young adult cohort from the HCP dataset, was applied to an independent ADNI dataset. We extracted activation maps from the last layer of the individualizing sub-model before converting them to probabilities. In these maps, a higher value for a vertex with respect to a parcel suggests a greater likelihood of belonging to that parcel. For each parcel, we calculated its “loading” by summing these values within and nearby the parcel. Linear regression was performed on normal participants to remove the influence of parcel size on loadings. The residuals from the regression, termed normalized loadings, are robust to parcel size and can be formulated as follows:

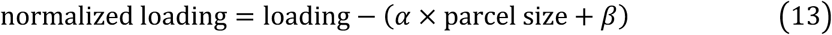

In contrast to morphological measures like volume or surface area, these loadings primarily reflect individual parcels’ relative shrinkage or expansion compared to their whole brain sizes, attributed to functional variations. To investigate the association between normalized parcel loadings and clinical groups (HC, MCI, and AD), generalized linear models (GLM) were employed. The clinical groups were treated as ordinal data (HC = 0, MCI = 1, and AD = 2), employing a Gaussian family with a logit link function. Covariates, including age, sex, head motion (measured by the average relative head motion during rsfMRI acquisition), and total intracranial volume, were controlled for in the analysis.

## 3. Results

### 3.1. Synthesizing task activation maps

In our evaluation of the task contrast map synthesis using the synthesis sub-model, we included 39 participants from the HCP test-retest dataset who underwent sMRI, rsfMRI, and tfMRI scans. We compared the synthesized maps with both group-averaged maps and individual target (i.e., test) and repeat (i.e., retest) maps. The group-average task contrast maps were created by averaging the maps from all participants in the HCP test dataset. As shown in Figure 2a, the synthesized maps exhibited the highest Pearson’s correlation coefficient (PCC) and Dice AUC with both the target and the repeat maps, outperforming the group-average maps. Detailed results for Dice curves and PCCs of each task contrast can be found in Supplementary Figure 2 and Supplementary Table 1, respectively. Figure 2b provides a visual representation of the Relational task Match contrast, featuring three randomly selected examples from the 39 participants. Echoing the overall findings, the synthesized maps in these examples demonstrated higher PCC and Dice AUC with both the test and retest maps compared to the group-average maps. Notably, the regions with high activation (marked by blue triangles) mirrored subject-specific patterns seen in the target and repeat measured maps, illustrating improved specificity over the group-average maps. Furthermore, the subject-specificity of the synthesized maps was evidenced by their higher similarity to each participant’s corresponding target maps rather than to the target maps of other participants (Supplementary Figure 3).

**Figure 2.**
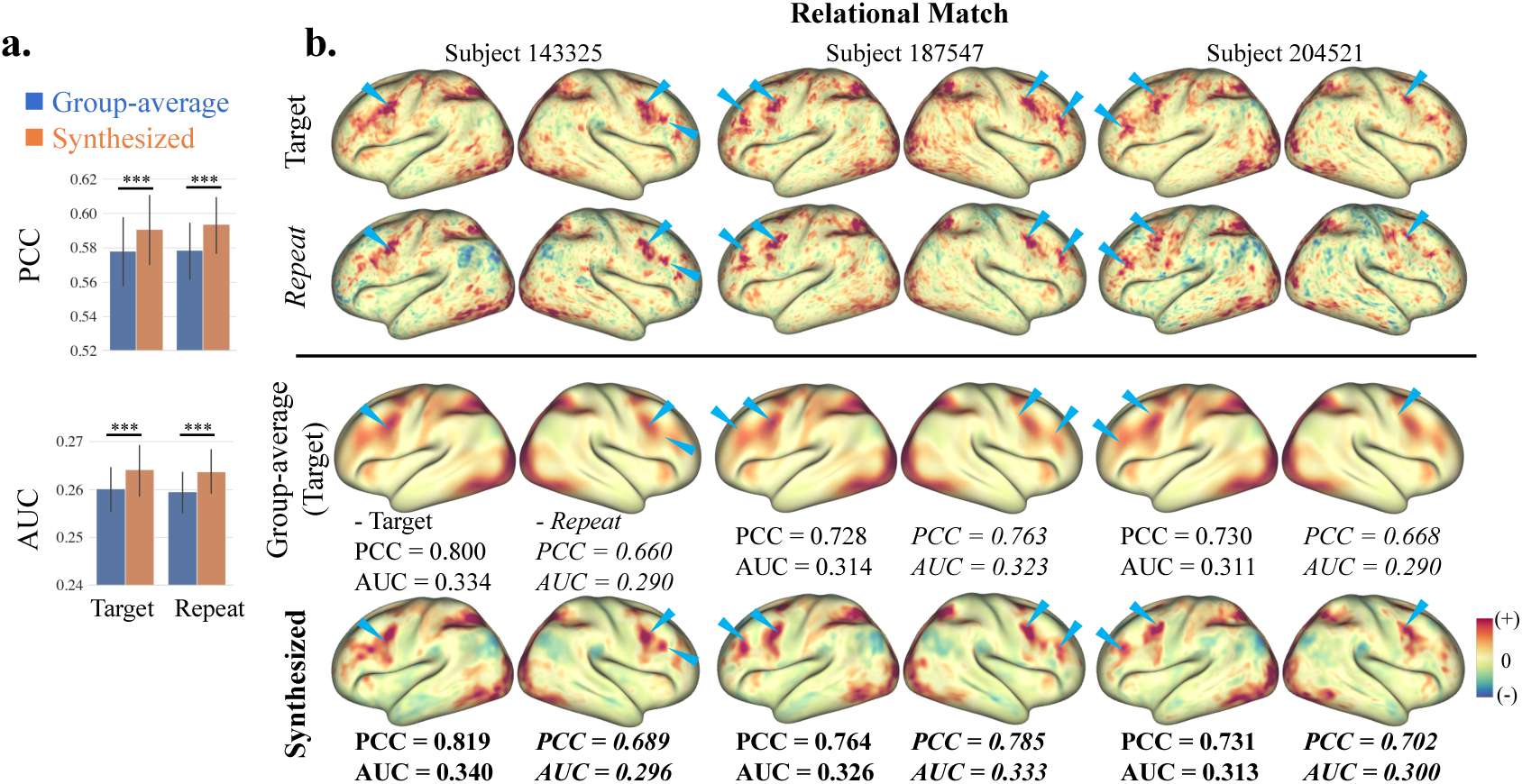
The synthesis contrast maps outperformed group-average contrast maps in capturing individual-specific features across HCP task contrasts. Quantitative metrics include Pearson correlation coefficient (PCC) and area under the curve (AUC) of Dice coefficient between target (i.e., test data) or repeat (i.e., retest data) and group-average or synthesized maps. **a.** The synthesis maps exhibit superior PCC and Dice AUC relative to group-average maps for both target and repeat maps among 39 test-retest participants across 24 task contrasts. The group-average maps were averaged from the target maps of the 39 participants. **b.** Target, repeat, group-average, and synthesized maps of Relational task Match contrast for three representative participants in the HCP dataset. Regions with high activations in synthesized maps showed subject-specific patterns similar to their target and repeat maps, marked by blue triangles. This suggests that TS-AI’s superior ability of capturing fine-grained, individual-specific features compared to group-average maps. For each participant, PCC and Dice AUC values for target and group-average/synthesized contrast maps are shown in normal font, and the values for repeat and group-average/synthesized contrast maps are shown in italics. ***, two-tailed paired t-test *p* < 0.001.

### 3.2. Test-retest reliability

We evaluated the reliability of individualized atlases using the HCP test-retest dataset (n = 39). As shown in Figure 3a, the effect size (Cohen’s d) of TS-AI (4.68) is higher than that of IMMP (3.59), suggesting better reproducibility and specificity of TS-AI compared to IMMP. Figure 3b presents individualized atlases for two participants chosen at random, derived from both the test and retest datasets using TS-AI and IMMP. In particular, for selected parcels across various lobes, a pronounced greater similarity is observable between test and retest atlases within individuals compared to the similarity across different individuals. Three additional TS-AI-derived individual atlases are also demonstrated in Supplementary Figure 4. These results highlight TS-AI’s ability to preserve intra-subject consistency while capturing inter-subject variability.

**Figure 3.**
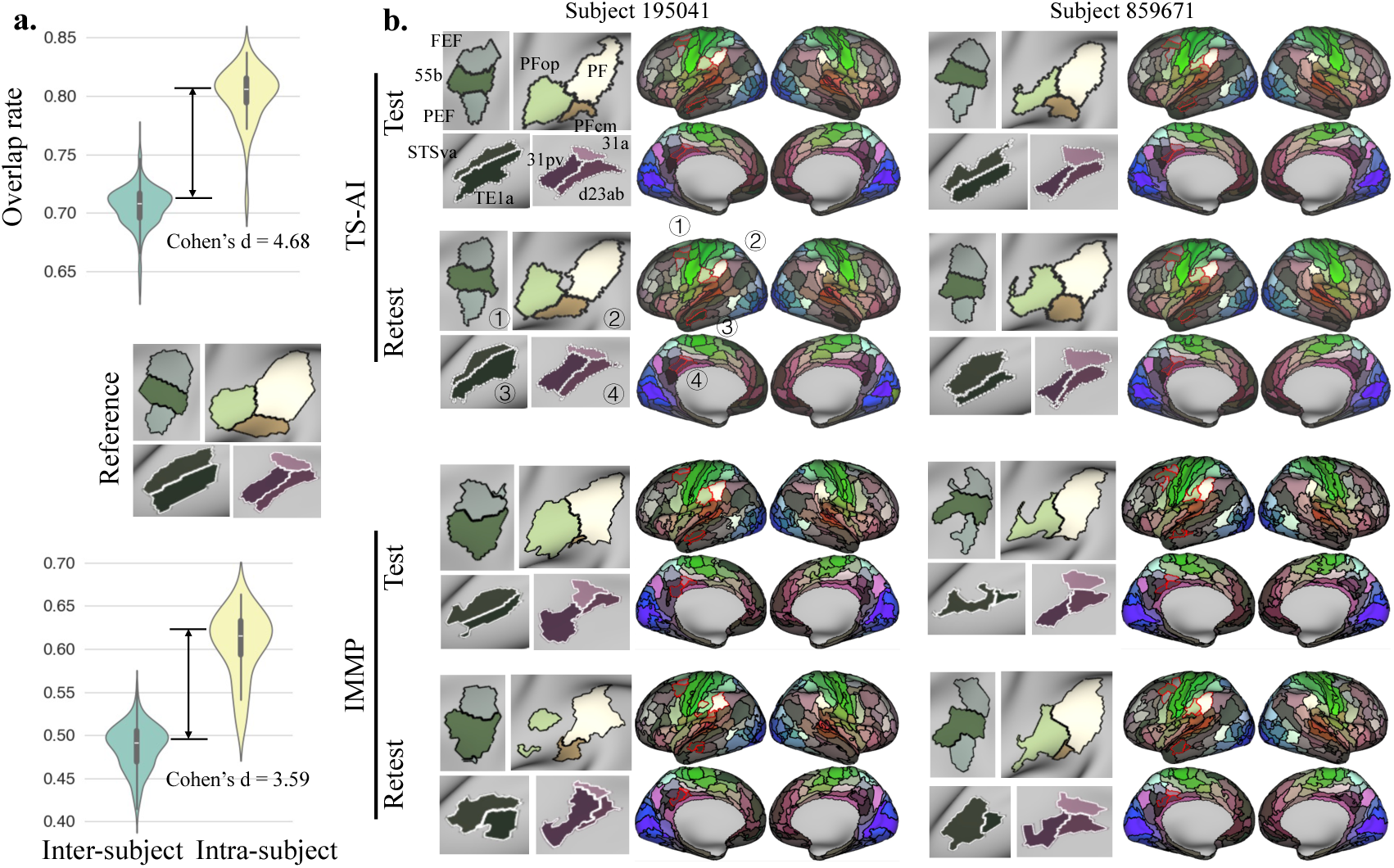
Reproducibility and specificity of TS-AI and IMMP using HCP test-retest dataset (n = 39). **a.** Violin plots show the overlap rates of inter- and intra-subject pairs, with higher effect size (Cohen’s d) of TS-AI than IMMP. **b.** Visual illustrations of TS-AI-based and IMMP-based individualized Glasser atlas of two randomly selected participants, with four magnified regional parcellations (red boundaries in whole-brain views) showing high inter-subject variability and intra-subject consistency. The middle panel between the two violin plots is the group-level Glasser atlas of the four exemplar regions. Visual comparisons for three additional participants are shown in Supplementary Figure 4. 31pv, area 31p ventral; 31a, area 31a; d23ab, area dorsal 23 a+b; FEF, frontal eye field; PEF, premotor eye field; PF, area PF complex; PFop, area PF opercular; PFcm, area PFcm; STSva, area STSv anterior; TE1a, area TE1 anterior.

### 3.3. Behavioral measures prediction

Kernel ridge regression (KRR) models were employed to predict 59 behavioral measures across six domains (alertness, cognition, emotion, motor, personality, and sensory) (Tian and Zalesky, 2018) using individualized Glasser parcellations derived from both IMMP and TS-AI. These predictions hinged on the similarity of parcellations among subjects. All available subjects (n = 953) were included in this predictive analysis. As shown in Figure 4a, TS-AI achieved a higher prediction performance (mean PCC = 0.113) compared to IMMP (mean PCC = 0.099), with this difference being statistically significant (two-sided paired t-test *p* < 0.05, FDR corrected) across all behavioral measures. Consequently, TS-AI demonstrated superior predictive performance over IMMP.

**Figure 4.**
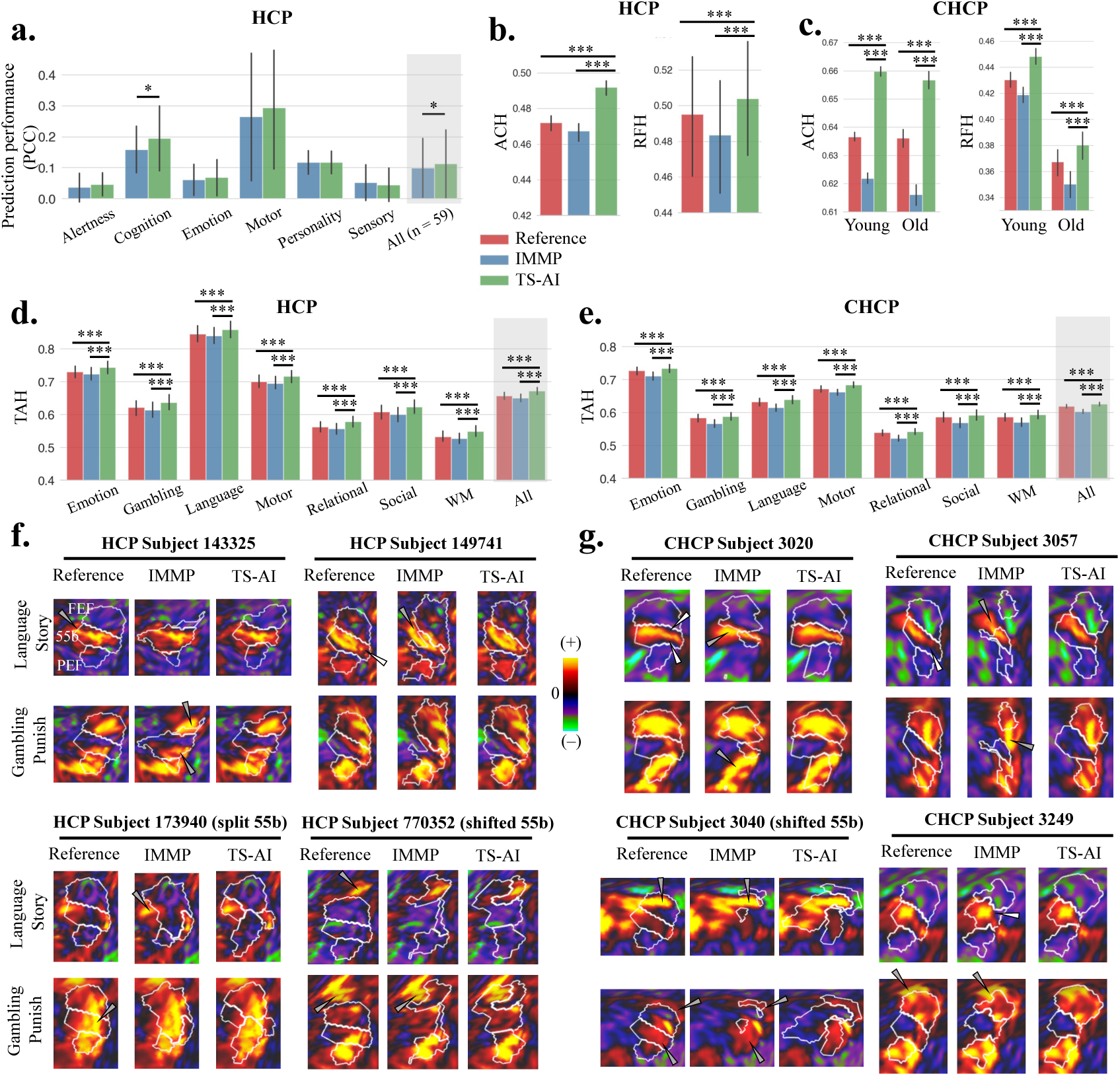
Validations of TS-AI using HCP and independent CHCP datasets. **a.** Kernel Ridge Regression models were employed to predict 59 behavioral measures across six domains using individualized parcellations based on the Glasser atlas (n = 953). Two-tailed paired t-tests were used to compare the performance of TS-AI and IMMP for each of the six subdomains and across all 59 behaviors. TS-AI showed significantly better performance than IMMP in the prediction of the Cognition subdomain and the aggregate of all measures. **b and c.** TS-AI demonstrated significantly higher anatomical connectivity homogeneity and resting-state functional homogeneity compared to the registration-based reference atlas and IMMP on HCP retest (b, n = 39) and CHCP young (c, n = 221, aged 18 ∼ 33 years) and old (c, n = 94, aged 48 ∼ 79 years) participants. **d and e.** TS-AI demonstrated significantly higher task activation homogeneity across all seven tasks on HCP retest (d, n = 39) and CHCP young (e, n = 160) participants. Note that only a part of CHCP young participants acquired tfMRI data. **f and g.** Task contrast maps of HCP retest (f) and CHCP (g) data for the Language task Story contrast and Gambling task Punish contrast in the left hemisphere were overlaid with the white contours of areas 55b, FEF, and PEF from three participants with atypical 55b (HCP subject 173940, HCP subject 770352, and CHCP subject 3040) and five other participants with typical 55b. The white arrows point to areas that were incorrectly included to a specific parcel, while the gray arrows point to areas that should have been included in the parcel but were omitted. TS-AI-derived parcel boundaries demonstrated better correspondence with the task contrast maps compared to IMMP-based and registration-based parcel boundaries. ACH, anatomical connectivity homogeneity; FEF, frontal eye field; IMMP, individualized multimodal parcellation (Glasser et al. 2016); PCC, Pearson’s correlation coefficient; PEF, premotor eye field; Reference, the reference atlas aligned to the individual space; RFH, resting-state functional homogeneity; TAH, task activation homogeneity; WM, working memory. Two-tailed paired t-tests were performed between TS-AI and IMMP/Reference with Bonferroni correction for (b) (c) and FDR correction for (a) (d) (e). *, *p* < 0.05,***, *p* < 0.001.

### 3.4. Homogeneity

We assessed the ACH, RFH, and TAH within individualized parcels. These assessments were calculated based on anatomical connectivity profiles, resting-state functional timeseries, and task contrast maps from the HCP retest dataset, respectively. Notably, these individualized parcellations were inferred from the HCP test dataset (n = 39). TS-AI demonstrated significantly higher ACH and RFH (Figure 4b), as well as TAH across seven tasks (Figure 4d) compared to the registration-based reference atlas and IMMP (two-sided paired t-test *p* < 0.001, Bonferroni corrected for ACH and RFH, and FDR corrected for TAH).

Previous evidence (Glasser et al., 2016) indicated that in the left hemisphere, area 55b, which is associated with language, is expected to coincide with regions of high activation during language tasks. Conversely, the frontal eye field (FEF) and premotor eye field (PEF), adjacent to area 55b, are implicated in saccadic eye movements and thus are typically activated during gambling-related tasks. As illustrated in Figure 4f, area 55b—individualized by TS-AI in four representative participants—demonstrated enhanced alignment with the region of intense activation for the Language task Story contrast, when compared to both the registration-based reference atlas and IMMP. Similarly, for the Gambling task Punish contrast, both FEF and PEF, as individualized by TS-AI, showed a more accurate alignment with the regions of intense activation. Moreover, TS-AI precisely identified the atypical topography of area 55b for Subjects 173940 and 770352, revealing split and shifted spatial arrangements that differ from its standard location in the group-level Glasser atlas.

### 3.5. Robust performances across datasets and atlases

To explore the generalizability of our model to independent datasets, we directly applied TS-AI and IMMP, both trained on the HCP first-fold dataset, to the independent CHCP dataset. The homogeneities of the individualized atlases in the CHCP dataset were assessed to reveal the generalization capability of TS-AI. Compared to the registration-based reference atlas and IMMP, TS-AI-derived individual parcellations obtained the highest ACH and RFH (two-sided paired t-test, p < 0.001, Bonferroni corrected) for both young and old groups of participants (Figure 4c). Since the old group did not acquire tfMRI data, the TAH was calculated only for the young group. In this case, TS-AI outperformed the registration-based reference atlas and IMMP (Figure 4e). Figure 4g features four representative subjects, in which areas 55b, FEF, and PEF individualized by TS-AI showed improved alignment with the regions of high task activation when compared to the registration-based reference atlas and IMMP.

Furthermore, to verify the robustness of our model across different atlases, we trained an individualizing sub-model using the Brainnetome atlas and evaluated its performance. The evaluation metrics included test-retest reliability, behavioral prediction, and homogeneities (Supplementary Figure 5). The effect size (Cohen’s d) between intra- and inter-subject overlap rates of the TS-AI-based individualized Brainnetome atlas was 4.81, confirming robust test-retest reliability. TS-AI outperformed IMMP on the average prediction performance of all behavioral measures (mean PCC = 0.099 vs. 0.089). Additionally, TS-AI significantly excelled in anatomical connectivity, resting-state functional connectivity, and task activation homogeneities compared to both the registration-based reference atlas and IMMP.

### 3.6. Interpretability via sensitivity analysis

To assess the sensitivity of the individualizing sub-model, a modality-wise sensitivity analysis was conducted, yielding sensitivity maps for each modality (Figure 5a). We observed that anatomical connectivity features contributed most to the individualization of the frontal and medial parietal regions. Task activation features showed a considerable impact on the temporal regions, while resting-state functional features were critical in individualizing the posteromedial cortices.

**Figure 5.**
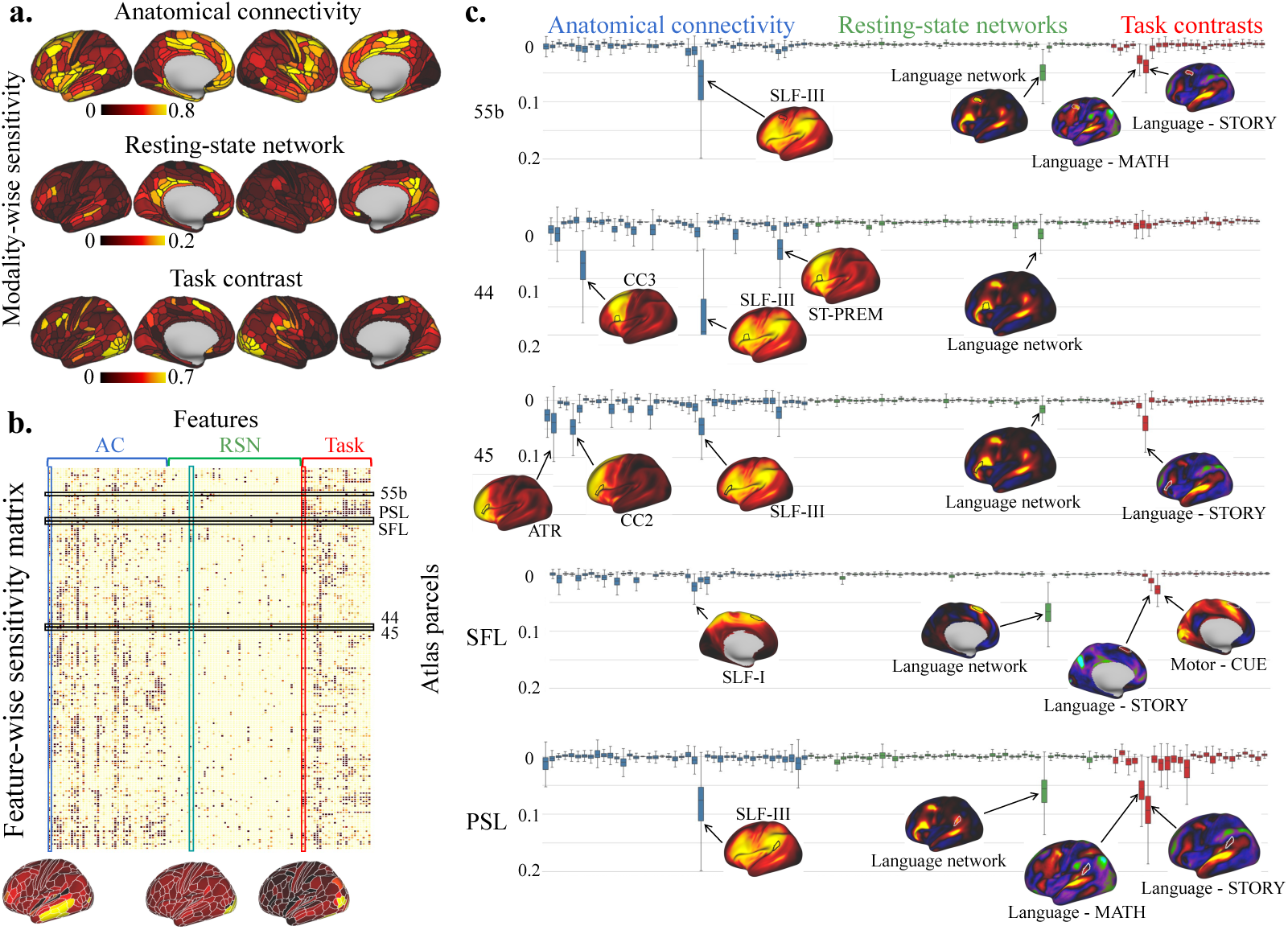
Sensitivity analyses of features contributing to the atlas individualizing sub-model. **a.** Modality-wise sensitivity map showing the contributing patterns of each modality to the atlas individualizing sub-model. Anatomical connectivity features exhibited high sensitivity in the lateral and medial frontal lobe, resting-state network features showed high sensitivity in posteromedial cortices, and task activation features contributed mostly to the occipital and temporal lobes. Bright yellow represents regions with highly discriminating features. **b.** The averaged feature-wise sensitivity matrix for the left hemisphere across subjects in the HCP first-fold test set. The sensitivity maps below illustrate three exemplar columns, each corresponding to the sensitivity patterns of distinct features. **c.** The boxplots depict five exemplar rows from (b), namely area 55b, area 44, area 45, superior frontal language area (SFL), and perisylvian language area (PSL), showcasing varying sensitivities for different parcels in the language network. On the bottom of the boxes, the corresponding average feature maps are overlaid with the boundaries of their respective parcels. AC, anatomical connectivity; RSN, resting-state network.

A feature-wise sensitivity analysis was performed to further explore the influence of specific input features. This analysis yielded a parcel-to-feature sensitivity matrix (Figure 5b), in which each element (𝑖, 𝑗) represents the sensitivity of parcel 𝑖 to feature 𝑗. When examining the contributions of language-related areas, including area 55b, area 44, area 45, perisylvian language area (PSL), and superior frontal language area (SFL), distinct patterns emerged, revealing how fiber tracts like the corpus callosum (CC) and the superior longitudinal fascicle (SLF) affect the individualization process (Figure 5c). Notably, area 44 is sensitive to the premotor part of CC (CC3) and area 45 to the genu part of CC (CC2). Considering the SLF, areas 55b, 44, 45, and PSL are sensitive to the ventral part of SLF (SLF-III), while area SFL is sensitive to the dorsal part of SLF (SLF-I). The locations of the mentioned segments of CC and SLF are presented in Supplementary Figure 6. The language network within resting-state functional networks plays a significant role in individualizing all selected language areas 55b, 44, 45, SFL, and PSL. Regarding task contrasts, areas 45 and SFL exhibited higher sensitivity with Language Story compared to Language Math tasks, areas 55b and PSL exhibited high sensitivity to both Language Story and Math tasks, while area 44 is neither sensitive to Story nor Math task.

We further conducted a correlation analysis between parcel-wise sensitivity and parcellation performance. First, we z-scored the feature-by-parcel sensitivity matrix (Figure 5b) across parcels for each feature and then averaged the sensitivity across features to obtain the mean normalized feature-wise sensitivity for each parcel (Supplementary Figure 7a). We then calculated the correlation between the mean normalized feature-wise sensitivity and the parcellation performance for each parcel, including ACH, RFH, TAH, and behavior prediction. Our results revealed that sensitivity is significantly correlated with ACH (r = 0.23, p < 0.001) and TAH (r = 0.16, p < 0.01), indicating that parcels with higher sensitivity exhibit high ACH and TAH (Supplementary Figure 7b). No significant correlations were found between sensitivity and RFH or behavioral prediction.

### 3.7. TS-AI applications in Alzheimer’s disease

The TS-AI model, trained on the HCP dataset, was applied to AD, MCI, and HC participants in the ADNI dataset. Compared to the registration-based reference atlas, TS-AI achieved significantly higher ACH and RFH for the AD, MCI, and HC groups (two-sided paired t-test, p < 0.001, Bonferroni corrected), as shown in Figure 6a. These results suggest that the TS-AI trained on HCP adults is effective for the ADNI dataset. We then investigated the variations in individual areal sizes identified by TS-AI across AD, MCI, and HC. GLMs were utilized to associate normalized parcel loadings with the clinical groups, controlling for covariates including age, sex, head motion, and total intracranial volume. Relatively faster shrinkage was observed in regions of bilateral medial temporal, cingulate, and lateral superior temporal areas, including structures such as the parahippocampus, entorhinal cortex, posterior area 24, and superior temporal sulcus dorsal regions (Figure 6c). In contrast, four regions displayed significantly slower shrinkage relative to the whole brain (Figure 6c). Detailed statistics of these significantly faster and slower shrinkage parcels are listed in Supplementary Table 4.

**Figure 6.**
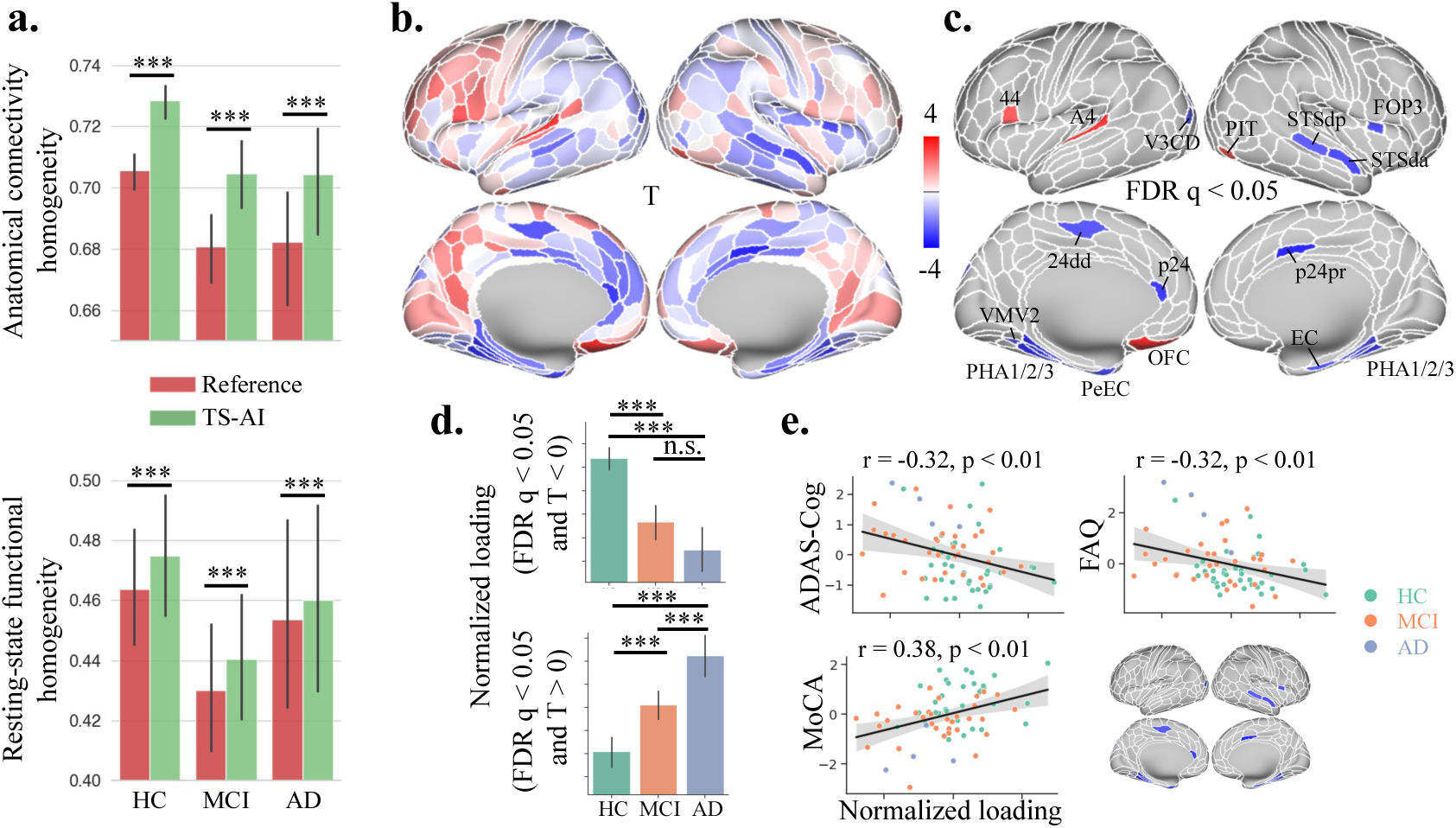
TS-AI applications in Alzheimer’s disease analyses. **a.** Compared to the registration-based reference atlas, TS-AI demonstrated significantly higher anatomical connectivity homogeneity and resting-state functional homogeneity for HC, MCI, and AD participants in ADNI dataset. **b.** The correlation between HC/MCI/AD groups and normalized parcel loadings assessed using generalized linear models, regressing out covariates including age, sex, head motion, and total intracranial volume. Blue and red parcels represent faster and slower shrinkage relative to the whole cortical surface from HC towards MCI and AD, respectively. The color bar indicates T values. **c.** The map shows the regions with significant changes from HC, MCI to AD (FDR q < 0.05). **d.** The total normalized loadings of significantly faster shrinkage parcels (FDR q < 0.05 and T < 0, blue) and slower shrinkage parcels (FDR q < 0.05 and T > 0, red) for HC, MCI, and AD. **e.** The normalized loadings in faster shrinkage parcels exhibit a significant negative correlation with ADAS-Cog and FAQ, and a significant positive correlation with MoCA. No significant correlation was observed between normalized loadings in slower shrinkage parcels and ADAS-Cog, FAQ, and MoCA. The y-axes of the scatter plots represent measures that were normalized and regressed out covariates including age, sex, head motion, and total intracranial volume. 24dd, dorsal area 24d; 44, area 44; A4, auditory 4 complex; EC, entorhinal cortex; FOP3, frontal opercular area 3; OFC, orbital frontal complex; p24, area posterior 24; p24pr, area posterior 24 prime; PeEc, perirhinal entorhinal cortex; PHA 1/2/3, parahippocampal area 1/2/3; PIT, posterior inferotemporal; STSda, area STSd anterior; STSdp, area STSd posterior; V3CD, area V3CD; VMV2, ventromedial visual area 2. ***, two-sample t-test Bonferroni corrected *p* < 0.001, n.s., not significant.

We hypothesized that the relatively faster and slower shrinkage of parcel sizes could be attributed to the progression of the disease. To test this hypothesis, we performed univariate regression analyses across the clinical groups, correlating the normalized loadings of regions exhibiting significantly faster or slower shrinkage with neuropsychological measures including Alzheimer’s Disease Assessment Scale-Cognitive Subscale (ADAS-Cog), Functional Activities Questionnaire (FAQ), and Montreal Cognitive Assessment (MoCA). Similar to the GLM analysis, we controlled for age, sex, head motion, and total intracranial volume. As shown in Figure 6e, the normalized loadings of significantly faster shrinkage parcels (T < 0 and FDR q < 0.05) were negatively correlated with ADAS-Cog (r = -0.32, *p* < 0.01) and FAQ (r = -0.32, *p* < 0.01), and positively correlated with MoCA (r = 0.38, *p* < 0.01). These trends are consistent with the changes of ADAS-Cog, FAQ, and MoCA during the progression of AD (Supplementary Table 5). No significant correlations were observed between the normalized loadings from significantly slower shrinkage parcels and neuropsychological measures.

### 3.8. Ablation study

An ablation study was performed to evaluate the impact of various input modalities on the atlas individualizing sub-model, as shown in Supplementary Table 6. The evaluation was performed based on Pearson’s correlation coefficient for behavior prediction, ACH, RFH, and TAH. To ensure a thorough assessment, we calculated the average ranking across these metrics for each combination of input modalities. The findings indicated that individualized atlases, which incorporated a combination of anatomical connectivity features, resting-state networks, and task contrast maps, achieved the highest overall performance.

## 4. Discussion

In the present study, we proposed the TS-AI method, an approach for cortical atlas individualization that is integrated with task contrasts synthesis using dMRI and rsfMRI. TS-AI first leveraged anatomical connectivity and resting-state functional networks to predict task contrast maps. Individualized atlases were then generated based on multimodal individual-specific features, including the predicted task contrast maps, as well as anatomical connectivity and resting-state functional networks. This stage focused on optimizing feature consistency within parcels while aligning the subject-specific parcellation with the reference atlas. Our results demonstrated that TS-AI achieved accurate and robust localization of parcels at the individual level. This was verified by more consistent anatomical connectivity, resting-state functional timeseries, and task-evoked activations within parcels, as well as improved prediction capacities for behavioral measures. Interpretability analysis further revealed that the model exhibited feature-specificity for different parcels, providing insights into the roles of various modalities in shaping individual functional regions. In addition, TS-AI identified accelerated shrinkage of medial temporal and cingulate regions during the progression of AD.

Contrast maps based on tfMRI are essential in delineating brain responses across subjects, and have been shown to accurately represent individual brain region topography (Glasser et al., 2016). The effectiveness of these modalities for synthesis was proved by previous studies, which indicated that anatomical connectivity (Saygin et al., 2012) and resting-state activation (Bernstein-Eliav and Tavor, 2022; Ngo et al., 2022; Tavor et al., 2016; Tik et al., 2023; Tobyne et al., 2018; Zheng et al., 2022) can partially reflect brain task activities. For instance, Saygin et al. (2012) used dMRI-measured anatomical connectivity to predict the functional activation of faces in the fusiform gyrus. Tavor et al. (2016) first suggested that resting-state functional connectivity can partially predict individual task-evoked responses using a set of regression-based models. Ngo et al. (2022) proposed a deep network model to predict task contrast maps from rsfMRI data. Given the challenge of acquiring a battery of tfMRI data in both research and clinical environments, TS-AI harnessed the benefits of the tfMRI during model training while allowing for its absence during inference. The ablation study highlighted the effectiveness of task contrast maps (Supplementary Table 6), showing that resting-state networks and tract-wise anatomical connectivity are not sufficient for generating individualized atlases and can be improved by the synthesized task contrast maps. The synthesized task contrasts serve as intermediate representations, supervised by ground truth task contrast maps. This is similar to deep supervision (i.e., auxiliary supervision) when considering TS-AI as an end-to-end model. Deep supervision involves introducing supervisory signals at the hidden layers of the network (Dou et al., 2016; Lee et al., 2015; Li et al., 2018). TS-AI learns to capture and encode relevant information from the connectivity features that are specifically useful for synthesizing task contrasts, which can ultimately benefit the individual parcellation.

In the TS-AI framework, both the task contrasts synthesizing and atlas individualizing sub-models receive anatomical connectivity and resting-state networks as input. Compared to deriving individual atlases solely from the synthesized task contrast maps, the reuse of resting-state networks and anatomical connectivity in Stage 2 improved behavior prediction, ACH, and RFH (Supplementary Table 6), demonstrating their contribution to individualized atlases. The repeated use of connectivity features in TS-AI is similar to the shortcut/skip connections in mainstream deep networks, such as ResNet (He et al., 2016), Transformer (Vaswani et al., 2017), and U-Net (Ronneberger et al., 2015). This architecture is considered efficient for enhancing the network’s representational ability and mitigating the risk of overfitting. We further compared TS-AI with a single-stage two-branch framework that synthesizes task activation and generates individualized atlases within a single model (see Supplementary Methods and Supplementary Figure 8 for a detailed description). The single-stage architecture was found to be inferior to the two-stage approach in terms of behavior prediction accuracy, ACH, RFH, and TAH (Supplementary Table 7), showing the effectiveness of the two-stage scheme. It is possibly because the two-stage framework employs separate decoders for synthesizing task contrasts and individualizing the atlas, while the single-stage framework shares a single decoder for both tasks, potentially leading to inferior representation capability of the model for individualization.

For a fair comparison between IMMP and TS-AI, we included resting-state functional networks and tract-wise anatomical connectivity as the input features for IMMP, which are identical to those used in TS-AI. Additionally, we compared IMMP with multimodal features identical to those in Glasser et al. (2016), including resting-state functional networks, ICA-based tfMRI independent components, as well as myelin and thickness maps (except visuotopic maps due to private processing). As shown in Supplementary Table 8, TS-AI outperformed both IMMP models in terms of behavior prediction accuracy, ACH, RFH and TAH. We also compared TS-AI with multimodal connectivity-based individual parcellation (MCIP), a non-deep learning method previously proposed by our research group (Cui et al., 2024). MCIP optimizes within-parcel homogeneity, spatial continuity, and similarity to a reference atlas using graph cut algorithm and can be directly applied to single subjects without training a model. In contrast, TS-AI, as a deep learning-based method, can automatedly capture complex links between brain features and subregional specialization. We found TS-AI (PCC = 0.113 ± 0.108) outperformed MCIP (PCC = 0.077 ± 0.071, both using a single rsfMRI session) in terms of behavior prediction, which is widely recognized as an important metric for evaluating individual parcellation, as it reflects the amount of individual-specific information encoded in individual atlases (Kong et al., 2019; Kong et al., 2021; Li et al., 2023; Li et al., 2019; Ma et al., 2022).

The lack of a definitive ground truth for individual atlas poses a challenge in the deep learning-based identification of individualized parcels. In the present study, we introduced a novel feature consistency loss function, designed to ensure the consistency of features within each parcel. The homogeneity of function and anatomy within parcels is well recognized and widely used in non-deep learning individualization methods (Chong et al., 2017; Honnorat et al., 2017; Kong et al., 2021; Parisot et al., 2017; Wang et al., 2015), therefore ensuring the credibility of the feature consistency loss as a prior assumption. Our evaluation of hyperparameter 𝜆 further clarified that the feature consistency loss improved the performance of the individualized atlas (Supplementary Table 3). Thus, the integration of feature consistency loss enables more complex deep networks to effectively capture spatial dependencies without being subject to overfitting risks.

Modality- and feature-wise sensitivity analysis captures the features that drive individual-specific cortical regionalization. The modality-wise sensitivity analysis revealed distinct spatial sensitivity patterns associated with different modalities. Specifically, anatomical connectivity features contributed most to the individualization of the frontal regions. In contrast, task activation features were more significant in the occipital and temporal regions, while resting-state functional features were most influential in the posteromedial cortices.

These findings support the notion that different modalities capture distinct individual regionalization differences, which aligns with previous research that emphasizes the complementary property of multimodal data in cortical parcellation (Glasser et al., 2016; Sui et al., 2014). Feature-wise sensitivity analysis further uncovered that specific regions were influenced by distinct features. For example, different segments of the CC notably impact various language areas: the premotor part of CC (CC3) for area 44 and the genu part of CC (CC2) for area 45. These observations align with previous findings linking CC to individual differences in language abilities (Bartha-Doering et al., 2021; Chiang et al., 2009; Dunst et al., 2014). However, unlike prior research which treated language-related areas as an entity, our study delineates the specific roles of different language areas in personal functional organization, each associated with various tract segments. In addition, the SLF is known to be relevant to language processing (Li et al., 2022), with its different segments suggested to serve varying functions in language (Gierhan, 2013). However, the specific areas within the language network that are relevant to these individual differences remain unknown. Our results demonstrated that the individual topographic differences in areas 55b, 44, 45, and PSL were closely linked to the ventral part of SLF (SLF-III), while area SFL was linked to the dorsal part of SLF (SLF-I). For resting-state functional networks, areas 55b, 44, 45, PSL, and SFL all showed particular sensitivity to the language network (Poologaindran et al., 2020; Rolls et al., 2022). Through sensitivity analyses, TS-AI offers insights into the region-specific basis that drives these individual differences in language abilities, as well as other primary and higher-order functions.

Our experimental findings provide compelling evidence that the sizes of the individual parcels exhibit associations with neuropsychiatric disorders. The normalized parcel loadings provided by TS-AI measure the relatively faster or slower shrinkage of individual parcels compared to the whole brain. Differing from morphological metrics like thickness, this variation primarily reflects impacts on functional anatomy. Specifically, we observed accelerated shrinkage in the parahippocampus, entorhinal cortex, cingulate cortex, and superior temporal regions during the progression of AD. These findings are consistent with previous studies that highlighted atrophy in the medial temporal lobe structures, particularly the parahippocampus (Cui et al., 2011; Ferreira et al., 2011) and entorhinal cortex (Cui et al., 2011), as critical predictors of the progression from pre-dementia stages to AD. In addition, altered interregional correlations, particularly in the medial temporal and medial frontal lobes, have been reported among MCI and AD groups (Yao et al., 2010). These observations suggest the potential of TS-AI in providing novel biomarkers for the diagnosis and prognosis of neuropsychiatric diseases. Hopefully, TS-AI could also hold promise for assisting in clinical applications, such as individual-specific region-targeting neuromodulation interventions like transcranial magnetic stimulation.

We acknowledge that the accuracy and effective resolution of cortical structural connectivity profiles estimated by fiber tractography may be susceptible to potential errors propagated from false positive and false negative fiber tractography caused by gyral bias, un-directional pathway trajectories, and underestimation of long-distance connections (Van Essen et al., 2014). Nevertheless, dMRI and tractography are indispensable in vivo techniques for estimating the anatomical connections of the whole human brain in a feasible manner, and fiber tractography can account for approximately 60% of the true neural fiber connections revealed by tracer studies (Donahue et al., 2016; Hayashi et al., 2021). In addition, the connectivity estimated from dMRI is more stable and state-independent, which has been used in the delineation of group-level parcellations for human (Fan et al., 2016), macaque (Lu et al., 2024), and marmoset (Liu et al., 2018) brains. Indeed, experimental results demonstrate that dMRI-based connectivity provides stable and complementary information to fMRI, and increases anatomical homogeneity as well as behavior prediction performance for individual parcellation (Supplementary Table 6).

In conclusion, we here present TS-AI, a novel multimodal deep learning approach for generating individual-specific brain atlases. TS-AI leverages dMRI and rsfMRI data to individualize an atlas, with the synthesis of various tfMRI contrasts. This methodology holds potential for precision medicine, particularly personalized diagnosis and treatment. In addition, TS-AI may offer valuable insights into how region-specific features contribute to individual differences in brain region topography.

## Data and code availability statement

The code for TS-AI is publicly available at https://github.com/YueCui-Labs/TS-AI and will be maintained and updated through this website.

All data used in this article is publicly available and the authors are not part of the consortia that have collected these data. Corresponding references where further information can be found, are made available in the manuscript.

## Acknowledgements

This work was supported by STI 2030 - Major Projects (No. 2021ZD0200402) and National Science Foundation of China (No. 82371486).

## Notes

### Competing Interest Statement

The authors have declared no competing interest.

